# Mitoxantrone modulates a glycosaminoglycan-spike complex to inhibit SARS-CoV-2 infection

**DOI:** 10.1101/2021.10.15.464595

**Authors:** Qi Zhang, Peter Radvak, Juhyung Lee, Yue Xu, Vivian Cao-Dao, Miao Xu, Wei Zheng, Catherine Z. Chen, Hang Xie, Yihong Ye

**Affiliations:** Laboratory of Molecular Biology, National Institute of Diabetes and Digestive and Kidney Diseases, National Institutes of Health, Bethesda, MD 20892, USA; Laboratory of Pediatric and Respiratory Viral Diseases, Division of Viral Products, Office of Vaccines Research and Review, Center for Biologics Evaluation and Research, United States Food and Drug Administration, Silver Spring, MD 20993, USA; National Center for Advancing Translational Sciences, National Institutes of Health, Rockville, MD 20850, USA

**Keywords:** SARS-CoV-2, COVID-19, viral entry, heparan sulfate proteoglycan/HSPG, glycosaminoglycan/GAG, spike, Mitoxantrone

## Abstract

Spike-mediated entry of SARS-CoV-2 into human airway epithelial cells is an attractive therapeutic target for COVID-19. In addition to protein receptors, the SARS-CoV-2 spike (S) protein also interacts with heparan sulfate, a negatively charged glycosaminoglycan (GAG) attached to certain membrane proteins on the cell surface. This interaction facilitates the engagement of spike with a downstream receptor to promote viral entry. Here, we show that Mitoxantrone, an FDA-approved topoisomerase inhibitor, targets a spike-GAG complex to compromise the fusogenic function of spike in viral entry. As a single agent, Mitoxantrone inhibits the infection of an authentic SARS-CoV-2 strain in a cell-based model and in human lung EpiAirway 3D tissues. Gene expression profiling supports the plasma membrane as a major target of Mitoxantrone but also underscores an undesired activity targeting nucleosome dynamics. We propose that Mitoxantrone analogs bearing similar GAG-binding activities but with reduced affinity for DNA topoisomerase may offer an alternative therapy to overcome breakthrough infections in the post-vaccine era.

---

The ongoing COVID-19 pandemic has claimed millions of lives worldwide and significantly damaged the global economy. Even with the development of multiple vaccines, the spread of SARS-CoV-2, the virus underlying the pandemic, is not completely halted due to breakthrough infections ^1–3^. As the hope for complete eradication of this devastating virus by vaccines dwindles, the call for effective therapies to save the lives of those infected, particularly the elderly and patients with pre-existing conditions has become urgent.

Once entering the human airway, SARS-CoV-2 invades the airway epithelial cells by receptor-mediated membrane fusion or endocytosis ^4–7^. Recent studies have identified several receptors mediating viral attachment and entry, which include angiotensin-converting enzyme 2 (ACE2), neuropilin-1, and tyrosine-protein kinase receptor UFO (AXL) ^8–11^. Among them, ACE2, as the key mediator of viral entry, has been extensively characterized ^12^. The N-terminal domain of ACE2 is composed of two lobes. SARS-CoV-2 uses the receptor-binding domain (RBD) in the spike (S) protein to contact the tip of one lobe. Neuropilin-1 was suggested recently as a distinct receptor for SARS-CoV-2 ^9,10^. This transmembrane receptor has two CUB (Complement C1r/C1s, Uegf, Bmp1) domains, one MAM (meprin/A5-protein/PTPmu) domain, and two coagulation factor domains. It interacts with the virus through a multi-basic site (after the cleavage of spike by a furin protease. The affinity of spike for neuropilin-1 *in vitro* is at micromolar levels, significantly weaker than its interaction with ACE2. AXL is another recently reported candidate receptor. It specifically interacts with the N-terminal domain of the S protein. Since AXL is not co-expressed with ACE2 in human lung tissues, it was suggested as an ACE2 independent receptor for SARS-CoV-2 ^11^.

The efficient entry of SARS-CoV-2 also requires two independent proteolytic reactions that activate the S protein. During viral assembly, the S protein is cleaved by a host furin protease, generating two segments: S1 and S2. S1 has the RBD domain mediating receptor binding ^7^. S2 drives membrane fusion as it contains a fusion peptide (FP, the functional fusogenic element) ^13,14^, which is activated by an additional cleavage at a site near the FP in the S2 segment. Depending on the subcellular location of the fusion reaction, the second cleavage could be mediated by either a cell surface protease (e.g. TMPRSS2) or a lysosomal protease such as Cathepsin L ^8,15^. In vitro, spike- and ACE2-mediated membrane fusion could even be activated by trypsin, a non-specific protease ^15^. Thus, the protease sensitivity of the S2 domain is a major determinant of viral entry efficiency ^4,7,16^.

Although viral receptors are generally considered as an ideal target for antiviral drug development, the existence of redundant SARS-CoV-2 receptors has imposed a challenge for this drug development strategy ^17–19^. We and others recently reported that SARS-CoV-2 uses the cell surface heparan sulfate proteoglycans (HSPGs) as an entry-assisting factor ^20,21^. HSPGs refers to a family of membrane proteins conjugated with negatively charged glycosaminoglycan (GAG) ^22^, which can attract a variety of cargos to the cell surface. Binding to GAG presumably increase the cargo dwell time and thus promote cargo-receptor interactions ^21,23^. Intriguingly, the polysaccharide of HSPGs can bind spike directly and even form a ternary complex with the spike RBD and ACE2 ^20,21^. It was proposed that the interaction of GAG with spike may facilitate spike activation. In this regard, drugs targeting the spike-GAG complex may interfere with the SARS-CoV-2 entry process independently of the viral entry receptors.

Our recent drug repurposing screen identified an FDA-approved anti-cancer drug named Mitoxantrone, which binds heparan sulfate (HS) and an HS analog (heparin) with high affinities ^21^. Mitoxantrone inhibits the entry of pseudo-viral particles coated with the SARS-CoV-2 S protein, but the underlying mechanism is unclear. Importantly, whether Mitoxantrone can similarly block the entry of authentic SARS-CoV-2 virus is unknown. In this study, we show that Mitoxantrone targets a spike-GAG complex, restricting the membrane fusion competency of spike. As a result, Mitoxantrone inhibits the entry of an authentic SARS-CoV-2 strain in human lung epithelial cells and in a human lung EpiAirway 3D tissue model. Thus, a small-molecule COVID-19 therapeutics could be potentially developed by modifying Mitoxantrone, removing the undesired topoisomerase binding activity while keeping the GAG binding moiety.

## Results

### Mitoxantrone targets a spike-GAG complex

Because heparin/HS binds to both Mitoxantrone and spike, we thought that heparin/HS might use the same binding site to interact with Mitoxantrone and spike. If so, Mitoxantrone could compete with spike for binding to HS on the cell surface, which would explain the observed antiviral activity. To test this idea, we conducted a pulldown experiment by incubating purified spike protein with heparin-conjugated beads in the absence or presence of excess Mitoxantrone. Surprisingly, immunoblotting showed that the presence of Mitoxantrone did not inhibit the spike-heparin interaction. Instead, it enhanced spike binding to heparin by ~7-fold. By contrast, Banoxantrone, a compound structurally homologous to Mitoxantrone, did not show any effect on the spike-heparin interaction (Figure 1A-C). These data suggest that the interaction between Mitoxantrone and GAG can stabilize a spike-heparin complex. This results in a ternary complex consisting of GAG, Mitoxantrone and spike.

**Figure 1.**
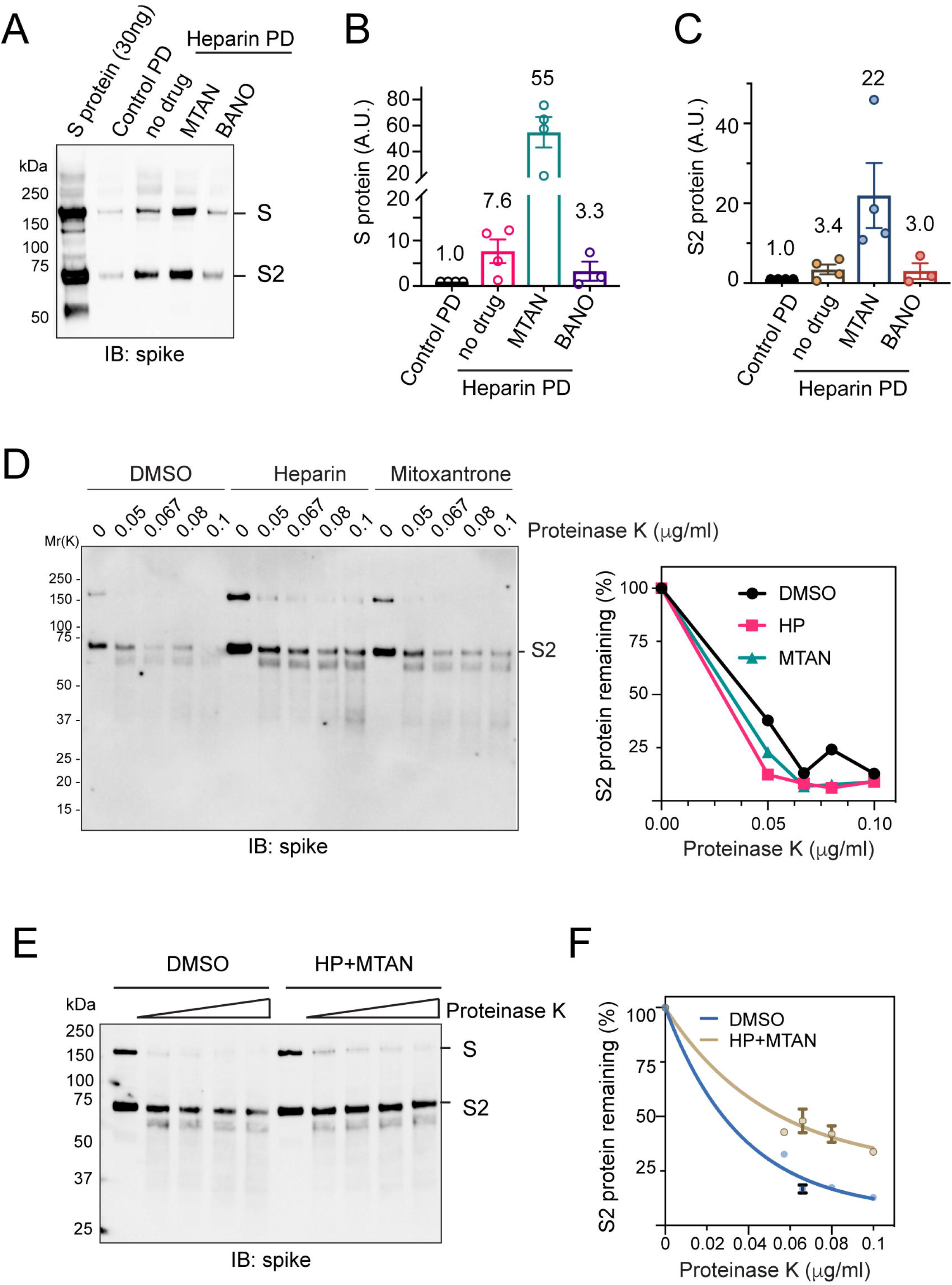
Mitoxantrone affects the spike-GAG interaction and alters the protease sensitivity of the spike protein. (A-C) Mitoxantrone but not Banoxantrone enhances the interaction of spike with heparin. (A) Heparin (HP) Sepharose beads were pre-incubated with a buffer in the absence or presence of 50 μM Mitoxantrone (MTAN) or Banoxantrone (BANO). After removing unbound drugs, we incubated the beads with recombinant spike (800ng). As a negative control, uncoated Sepharose beads were used. Proteins precipitated were analyzed by immunoblotting with spike antibodies. (B and C) Quantification of the full-length spike (S) or the furin-cleaved spike (S2) in A. n=4 independent experiments. (D) Either heparin or Mitoxantrone alone does not affect protease sensitivity of spike. Spike protein incubated with heparin (25 μM) or mitoxantrone (25 μM) or with a buffer control was treated with an increased concentration of proteinase K for 10 min. After quenching the protease, samples were analyzed by immunoblotting with spike antibodies. (E, F) GAG binding in the presence of Mitoxantrone increases the protease resistance of spike. As in D, except that spike was treated with heparin and Mitoxantrone together before proteolysis. (F) Quantification of the S2 band in E. n=3 independent experiments. Error bars, S.E.M.

The increased affinity between spike and heparin by Mitoxantrone suggests that the conformational dynamics of spike may be changed in the ternary complex. To test this hypothesis, we treated purified spike with low concentrations of proteinase K in the absence or presence of Mitoxantrone and/or heparin. Both the full-length spike and the furin-processed S2 domain were readily degraded by proteinase K in the absence of heparin and Mitoxantrone or in the presence of either of them (Figure 1D). However, when Mitoxantrone and heparin were both present, the S2 domain was more resistant to proteolysis, suggesting that in the ternary complex, spike is in a state less prone to activation by protease (Figure 1E, F).

### Mitoxantrone inhibits spike-mediated membrane fusion

The reduced sensitivity of the S2 domain to protease by heparin and Mitoxantrone treatment raised the possibility that Mitoxantrone may block spike-mediated membrane fusion. To test this, we established an *in vitro* membrane fusion assay by mixing two populations of HEK293T cells expressing spike and ACE2-GFP, respectively (Figure 2A). To identify spike-positive cells, the spike expressing plasmid was co-transfected with a mCherry-expressing plasmid. When mCherry and spike co-transfected cells were incubated with GFP-transfected cells or when mCherry-expressing cells were mixed with ACE2-GFP positive cells, no cell fusion was observed because green and red cells were segregated (Figure 2B, panels 1, 2). However, when spike-, mCherry-positive cells are incubated with ACE2-GFP cells at 1:1 ratio for 90 min, we observed rapid cell-cell fusion, resulting in the mixing of green and red fluorescence in giant syncytia filled with multiple nuclei (panels 3, 5). Interestingly, when cell fusion was carried out in the presence of Mitoxantrone, the cell-cell fusion was significantly inhibited as indicated by increased number of unfused cells or reduced syncytium size (panel 4, quantification in Figure 2C). Altogether, we concluded that Mitoxantrone modulates a GAG-spike complex, changing the protease sensitivity of the S2 domain, which inhibits spike-mediated membrane fusion.

**Figure 2.**
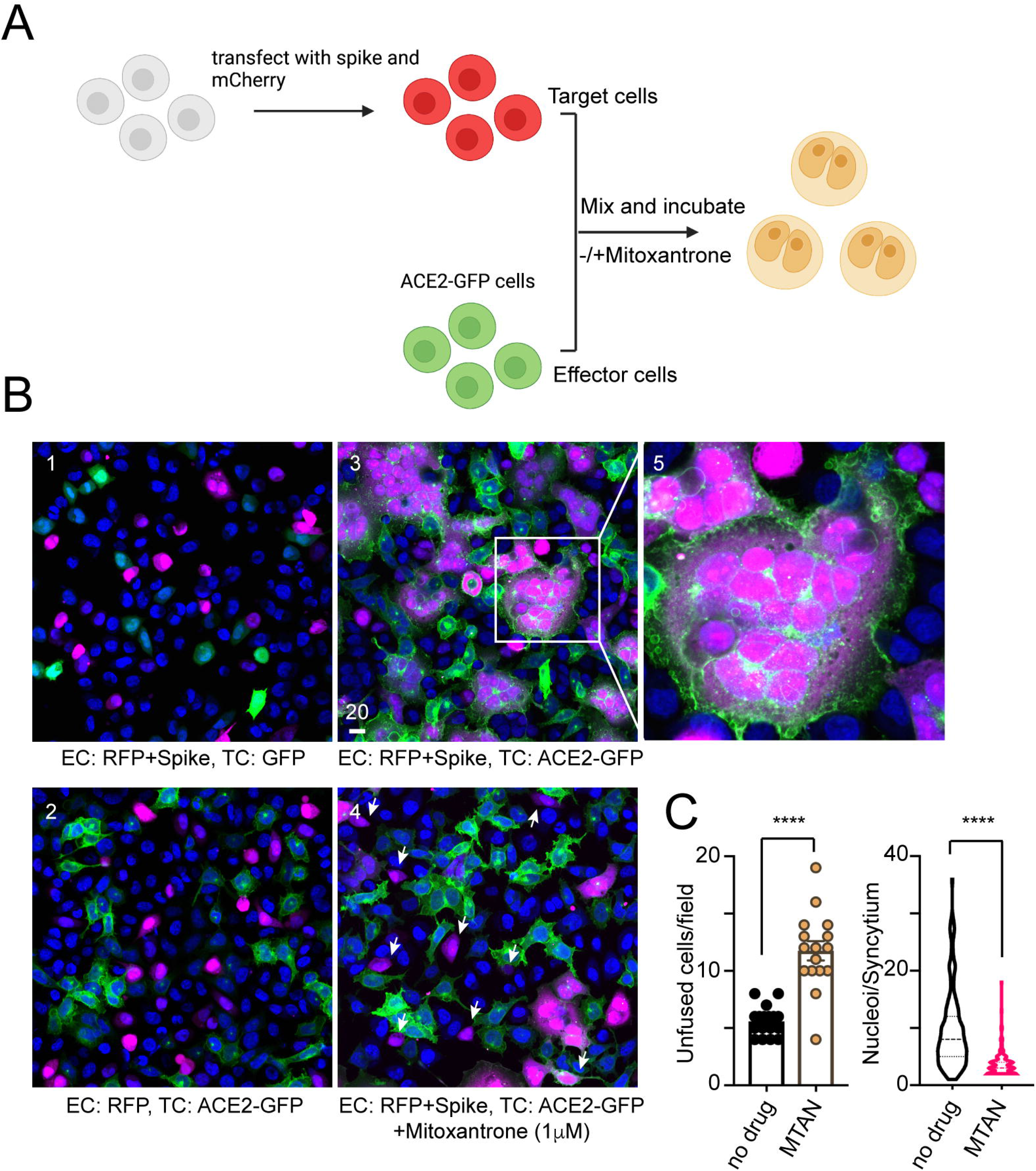
Mitoxantrone inhibits spike- and ACE2-mediated membrane fusion. (A) A scheme of the membrane fusion assay. (Created by Biorender.com) (B) Mitoxantrone inhibits spike and ACE2-mediated membrane fusion. Effect cells (EC) and target cells (TC) transfected with the indicated plasmids were mixed in equal numbers and cultured in the absence (panels 1-3) or presence (panel 4) of Mitoxantrone (1 μM) for 90min. Cells were fixed and imaged by confocal microscopy. Panel 5 shows an enlarged view of the boxed area in panel 3. Arrows in panel 4 show example of unfused target cells. Scale bar, 20 μm. (C) Quantification of the experiments shown in B. The left panel shows the number of unfused cells/field, while the right panel shows the number of nuclei/syncytium. ****, p<0.0001 by unpaired student’s t-test. n=2 independent experiments.

### Mitoxantrone inhibits the cell entry of an authentic SARS-CoV-2 strain

To examine whether Mitoxantrone could block the entry of an authentic SARS-CoV-2 strain, we treated cells with Mitoxantrone or as a control, with DMSO, and then exposed these cells to SARS2-CoV-2 (USA-WA1/2020) at an MOI of 0.1 for 5 hours. We then stained the cells with an antibody specific for the S protein to detect viral particles inside the cells. To demonstrate the antibody specificity, uninfected cells were stained in parallel, which showed only a low diffusive background signal (Figure 3A, top panels). By contrast, control cells infected with SARS-CoV-2 contained many bright spike-positive puncta throughout the cytoplasm (Figure 3A, middle panels). Strikingly, when Mitoxantrone-treated cells were infected and stained, we detected a significantly lower number of spike-positive signals compared to DMSO-treated cells (Figure 3A, bottom panels; Figure 3B). These results suggest that like pseudo-viral particles, the infection of authentic SARS-CoV-2 is also inhibited by Mitoxantrone.

**Figure 3.**
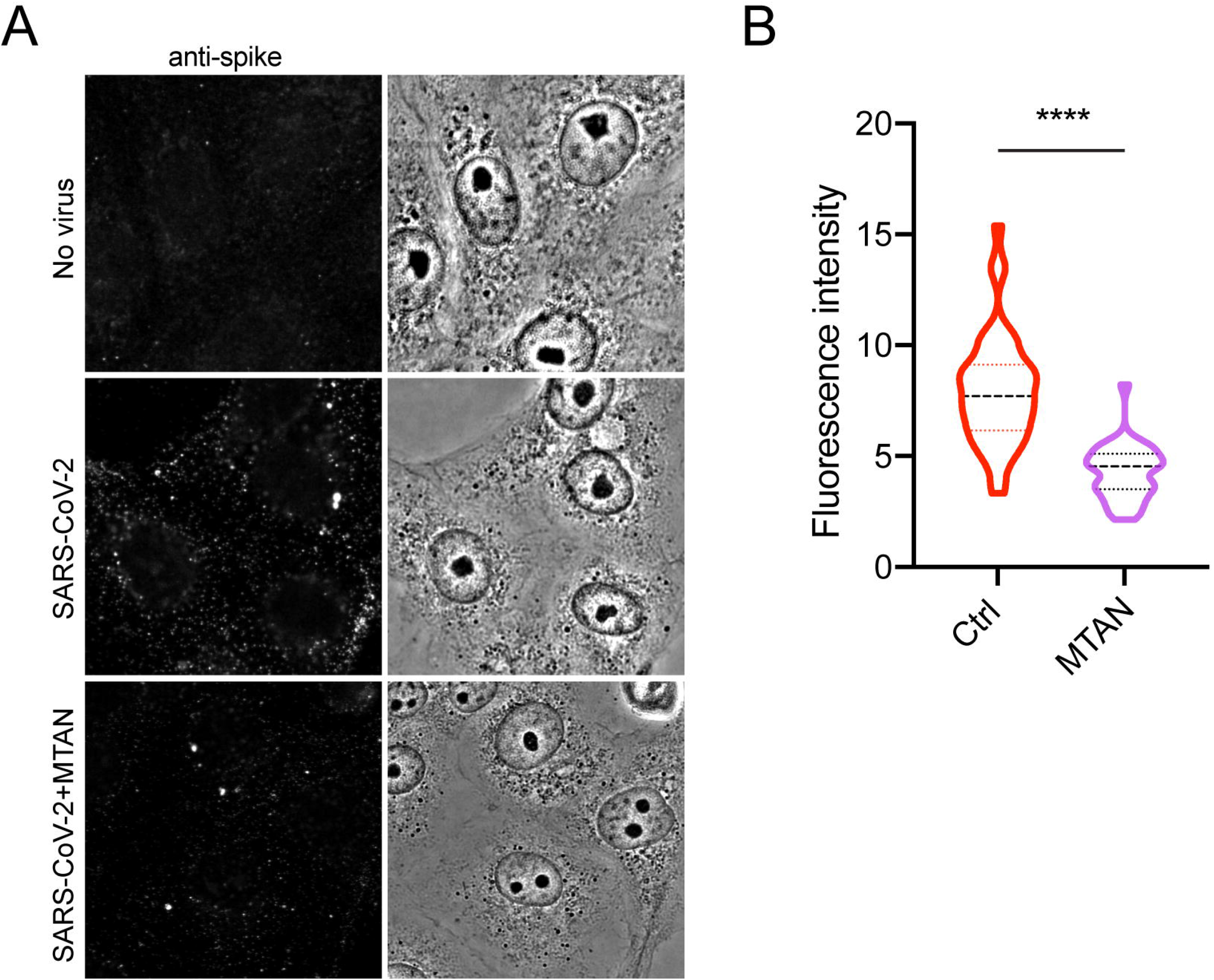
Mitoxantrone inhibits the entry of SARS-CoV-2 into a lung epithelial cell line. (A) Cells grown in monolayer were treated with Mitoxantrone (200nM) and then infected with SARS-CoV-2 (USA-WA1/2020) at an MOI of 0.1. Five hours later, cells were fixed and stained with rabbit anti-spike antibodies in combination with goat anti-rabbit IgG conjugated with Alexa Fluor 488 (left panels). (B) Quantification of spike fluorescence intensity in A. ****, p<0.0001 by unpaired student’s t-test, n=3 independent experiments.

### Mitoxantrone inhibits SARS-CoV-2 infection in a human 3D EpiAirway model

We next tested the anti-SARS-CoV-2 activity of Mitoxantrone in a model that mimics viral infection in the human airway. To this end, we chose human lung epithelial cell-derived 3D EpiAirway tissues, which contain three pseudostratified layers including an apical ciliated surface, the underlying mucociliary epithelium and a basal membrane. The tissues were cultured at the air-liquid interface (ALI) and can be infected with SARS-CoV-2 from the apical side (Figure 4A) ^24,25^. One hour before the infection, Mitoxantrone of different concentrations, or as a negative control, DMSO, was added to the medium. For positive control, we treated the tissues with Remdesivir, a nucleoside analog known to inhibit SARS-CoV-2 replication ^26^. After infection, uninfected viral particles were removed by gentle wash. Tissues were maintained in test article-containing medium on the basolateral side for 24 or 96 hrs. The tissues were carefully washed from the apical side. The levels of infection-competent SARS-CoV-2 virions in the washes were determined. As expected, Remdesivir treatment (2 μM) dramatically reduced the viral load in the wash collected at 24 hours post-infection but only partially reduced the virus titer in the wash collected at 96 hrs post-infection. By contrast, the DNA damaging drug Bleomycin had no such effect (Figure 4B). These results confirm the anti-SARS-CoV-2 activity of Remdesivir, which blocks viral replication to reduce the viral load. Consistent with our observation in cultured cells, Mitoxantrone treatment also dose-dependently reduced the viral load in the washes at both 24 and 96 hrs post-infection (Figure 4C, D). Compared to Remdesivir, the anti-SARS-CoV-2 effect of Mitoxantrone is much more potent as it achieved a similar level of inhibition at a concentration 20-fold lower than Remdesivir.

**Figure 4.**
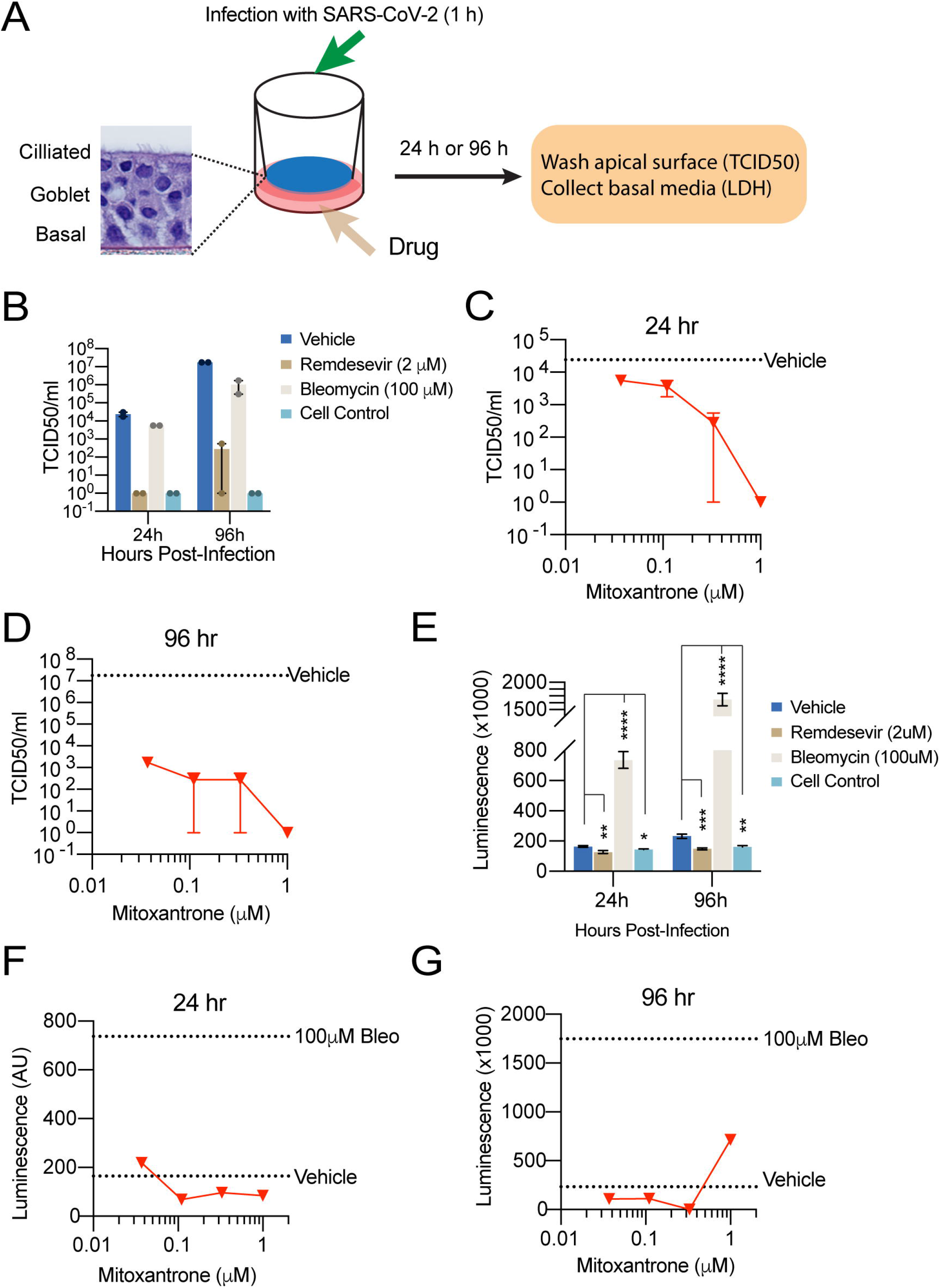
Mitoxantrone inhibits SARS-CoV-2 infection in an EpiAirway 3D tissue model. (A) A schematic diagram of the experimental design. (B) Remdesivir (2 μM) but not Bleomycin (100 μM) inhibits SARS-CoV-2 infection. 24 or 96 hrs after drug treatment and viral infection (MOI 0.1), the cell surface was washed. The viral titer (TCID50) in the wash was determined. (C, D) Mitoxantrone inhibits SARS-CoV-2 infection in the EpiAirway 3D model. TCID50 was determined either 24 hrs (C) or 96 hrs (D) after the organoids were treated with the drug at the indicated concentrations and then air-infected with SARS-CoV-2 at MOI 0.1 for 1 hr. The cells were washed from the apical side to remove the virus in the cell exterior and then incubated for 24 (D) or 96 hrs (E). Cells were rinsed from the apical side again and viral titers in the wash were determined. The dashed lines indicate the viral titer from cells infected without Mitoxantrone. (E) Bleomycin (Bleo. 100 μM) but not Remdesivir (2 μM) induces cell death, releasing LDH as determined by a luciferase assay. *, p<0.05, **, p<0.01, ***, p<0.001, ****, p<0.0001 by unpaired student’s t-test. n=2 tissues per test, each with 3 technical repeats. (F, G) Mitoxantrone inhibits SARS-CoV-2-induced cytotoxicity in the EpiAirway 3D model. As in D and E except that the concentration of LDH in the wash was determined.

To further confirm the anti-SARS-CoV-2 activity of Mitoxantrone, we measured the level of lactate dehydrogenase (LDH) in the basal culture media. LDH is a cytosolic enzyme released to the cell exterior upon cell death. Accordingly, high levels of LDH in the basal media from tissues treated with the apoptosis-inducing drug Bleomycin were detected (Figure 4E). A small increase in LDH was also observed in the basal media from SARS-CoV-2-infected cells compared to uninfected control, which was prevented if cells were pretreated with Remdesivir (Figure 4E). Virus-induced LDH release was also inhibited in Mitoxantrone-treated tissues at 24 hrs (Figure 4F). At 96 hrs post-infection, a low concentration of Mitoxantrone continued to lower the LDH level in the basal media compared to vehicle-treated control (Figure 4G). However, we observed a small increase in LDH level in 1 μM Mitoxantrone-treated samples (Figure 4G). This is probably due to Mitoxantrone’s intrinsic cytotoxicity (see below). Collectively, these data suggest that Mitoxantrone is a potent anti-SARS-CoV-2 entry inhibitor.

### Gene expression profiling reveals an undesired activity of Mitoxantrone

To understand the global impact of Mitoxantrone treatment on cells and to identify target inhibition that is irrelevant to viral entry, we analyzed the mRNA sequencing profile of HEK293T cells exposed to Mitoxantrone. This treatment dramatically changed the expression of many genes (Figure 5A). Gene ontology (GO) analysis of the 828 genes upregulated by Mitoxantrone by at least 2-fold with adjusted p-values less than 0.05 in triplicated experiments showed that the enriched processes fall into two categories: they are either associated with nucleosome assembly or disassembly or linked to events at the plasma membranes such as cellular response to stimulus, response to nutrient levels, regulation of cell-cell adhesion, etc. (Figure 5B). The latter is consistent with the observed interaction of Mitoxantrone with the cell surface GAG, which could induce compensatory gene expression changes to maintain proper cellular responses to extracellular stimuli. On the other hand, the upregulation of histone genes and genes regulating nucleosome assembly (Figure 5C) is consistent with the reported interaction of Mitoxantrone with nucleic acid and DNA topoisomerase (Figure 5D), which is thought to account for the cytotoxicity of Mitoxantrone in cancer therapy. Structural analyses show that the dihydroxy-anthraquinone moiety of Mitoxantrone forms extensive hydrophobic interactions with the DNA/topoisomerase complex (Figure 5E, F). Since the anti-SARS-CoV-2 activity has been linked to the two symmetric arms, we proposed that modification of the anthraquinone moiety may reduce cytotoxicity while maintaining the GAG-binding and thus the anti-SARS-CoV-2 function of Mitoxantrone.

**Figure 5.**
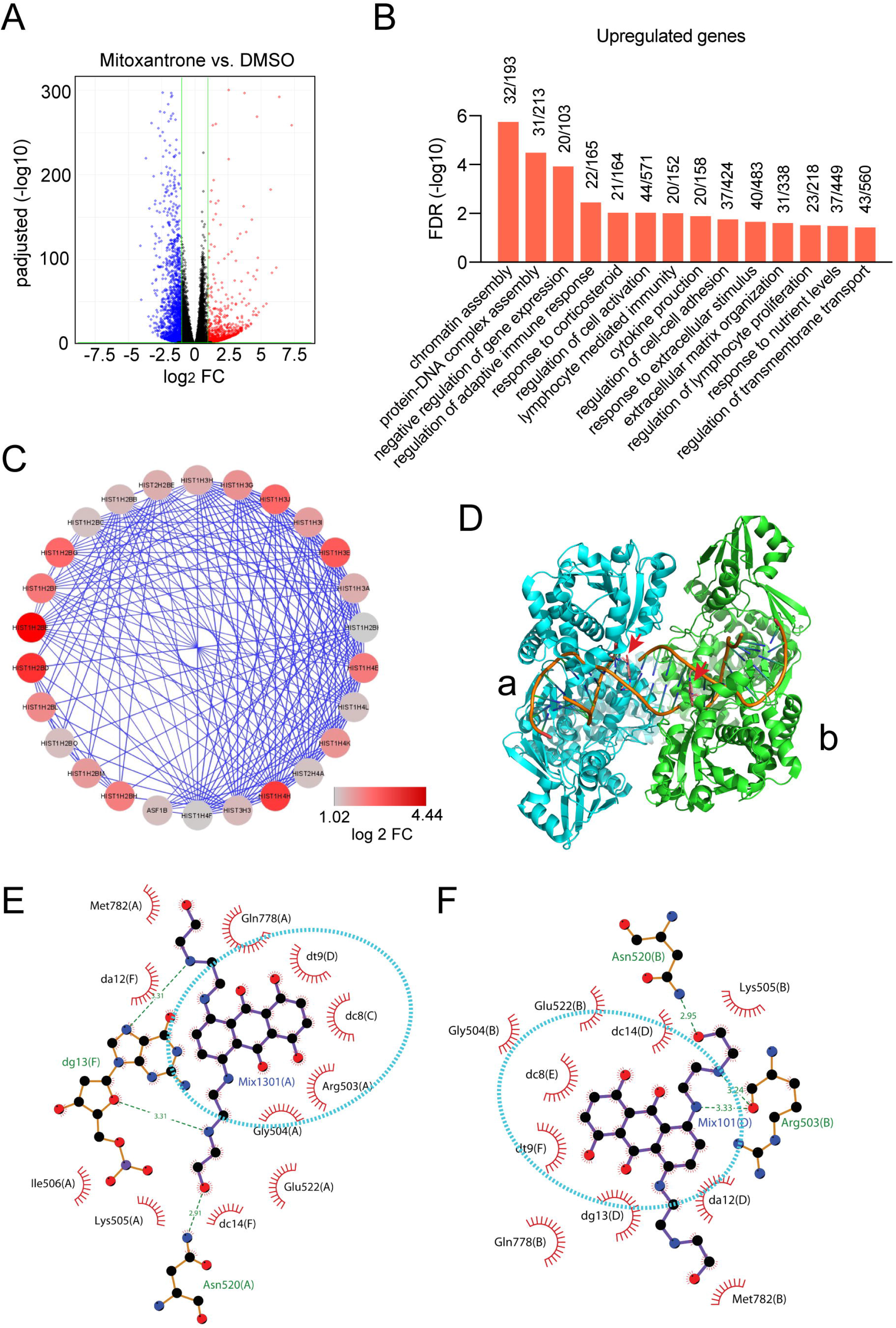
Gene expression profiling shows two major cellular targets of Mitoxantrone. (A) A volcano plot showing genes upregulated or downregulated by Mitoxantrone in HEK293T cells. (B) A list of significant pathways upregulated by Mitoxantrone. (C) Genes encoding nucleosome proteins are highly upregulated by Mitoxantrone. Shown is a gene interaction network highlighting nucleosome assembly-related genes induced by Mitoxantrone. Colors indicate fold change. (D) A crystal structure shows the binding of Mitoxantrone to a DNA-topoisomerase complex. Arrows indicate the two Mitoxantrone molecules each associated with a topoisomerase monomer. (E, F) A LigPlot view of the molecular interactions between Mitoxantrone and the DNA-topoisomerase complex.

## Discussion

The rapid development and deployment of COVID-19 vaccines have significantly slowed down the current pandemic. Nevertheless, the evolution of new viral strains combined with the disparity in global vaccine distribution has led to new waves of infected cases and deaths. Thus, an effective and economical therapeutic agent is urgently needed, particularly in developing countries that have limited access to vaccines. Our study suggests that Mitoxantrone derivatives with reduced affinity to DNA-topoisomerase complex may be a promising COVID drug candidate.

Mitoxantrone has been approved for treating acute myeloid lymphoma, prostate cancer, and multiple sclerosis. The anti-cancer activity has been attributed to its interaction with a DNA-topoisomerase complex, which inhibits DNA replication ^27^. Our study suggests that Mitoxantrone also has a high affinity for heparan sulfate, and this activity can be separated from its toxicity ^21^. Consistent with the newly established role of GAG in ACE2-dependent entry of SARS-CoV-2 ^20^, Mitoxantrone effectively inhibits the infection by spike-pseudotyped viral particles ^21^. Our new findings now suggest a mechanism by which Mitoxantrone interferes with spike-mediated viral entry. The binding of Mitoxantrone to GAG does not prevent its association with spike. Instead, it stabilizes spike in a complex with GAG, increasing its resistance to protease digestion. Because the conversion of spike into a fusion competent state requires a proteolytic clip downstream of furin-mediated cleavage, the presence of Mitoxantrone in a GAG-spike complex inhibits the fusogenic activity of spike, which in turn diminishes the infectivity of SARS-CoV-2. The binding of Mitoxantrone to the GAG-spike complex also interferes with ACE2-dependent endocytosis of viral particles ^21^, probably by altering a spike-ACE2 interaction. Thus, Mitoxantrone has at least two targets in mammalian cells. A straightforward strategy for developing Mitoxantrone into an antiviral agent is to generate derivatives that only bind to GAG with high affinities.

Compared to other viral entry inhibitors under development, Mitoxantrone is unique as it is a small molecule targeting a host factor required for viral entry. Thus, it has several advantages. For example, several neutralizing antibodies have been approved for use in patients with mild COVID symptoms. Many studies also reported promising anti-SARS-CoV-2 activities for nanobodies against spike in animal models ^28–32^. Although antibody-based therapies are generally safe and effective, they requires high doses, which make these therapies quite expensive. Additionally, most neutralizing antibodies recognize the RBD of spike, which is rapidly evolving as the virus continues to spread. Consequently, new variants with mutated spike may not be recognized by existing antibodies. In this regard, a safe, small molecule inhibitor targeting a host factor required for viral entry may offer an alternative therapeutic option that can be used either as a single agent or in combination with spike antibodies and other antiviral agents.

Drugs targeting host factors have been successfully developed to treat other infectious diseases. For example, Maraviroc, an anti-HIV agent targeting the host membrane protein CCR5 has been approved for AIDS treatment ^33^. Therefore, it is tempting to generate drugs with similar GAG-binding activities as Mitoxantrone but improved safety profiles. Our structural analyses suggest that the two hydroxyl groups in the anthraquinone moiety of Mitoxantrone may be the best place to introduce modifications, which are expected to reduce its binding to DNA-topoisomerase. Given the broad role of the cell surface GAGs in pathogen entry, Mitoxantrone derivatives may even be used to treat a new viral infection as long as the virus exploits the cell surface GAG for attachment and entry.

## Materials and Methods

### Reagents and plasmids

Spike protein was purchased from Sino Biologicals. Mitoxantrone was purchased from Microsource (Cat #01503278) and stored by the NCATS compound management department. Banoxantrone was purchased from Sigma (Catalog number: SML1854). pcDNA3.1-SARS-CoV-2-Spike plasmid was obtained from BEI resource ^7^. pLV-Mcherry was obtained from Addgene (Catalog number: 36084). Hoechst 33342 staining solution and proteinase K were purchased from Thermo Fisher Scientific (Catalog number: 62249 and AM2546).

### Heparin sepharose beads pulldown and protease K digestion

Heparin beads pulldown assays were performed as previously reported ^21^. For protease treatment experiment, 1.28 μg spike protein was incubated with either DMSO, Mitoxantrone (25 μM), heparin (25 μM) or Mitoxantrone together with heparin in the presence of 0.1 M of Tris-HCl (pH7.4) in pure water (220 μl of the total volume). Mitoxantrone and heparin were pre-mixed before adding to the S protein to avoid unwanted precipitations. After incubation at room temperature for 15 min, the reactants were aliquoted into 40 μl per tube. 10 μl pre-diluted proteinase K was added to make the final concentrations as 0.05, 0.067, 0.08, or 0.1 μg/ml. The reactions were further incubated for 5 min at room temperature and then quenched by preheated 4x Laemmli buffer and heating. The samples were resolved by SDS-PAGE electrophoresis followed by immunoblotting.

### Spike- and ACE2-mediated cell fusion assay

HEK293T target cells expressing spike protein and mCherry were generated by transfecting cells with pcDNA3.1-SARS-CoV-2-Spike and pLV-Mcherry at 1: 1 ratio. Effect cells were HEK293T cells stably expressing ACE2-GFP, as described previously ^21^, or as a control, HEK293T cells transfected with pEGFP. To initiate cell-cell fusion, effect cells and target cells were mixed at 1:1 ratio in a live cell imaging chamber and placed in the incubator for 90 min. Cells were then fixed by 4% paraformaldehyde in phosphate buffer saline and then stained with a Hoechst 33342 staining solution to label nuclei. Cell were imaged by a Zeiss LSM780 confocal microscopy.

### SARS-CoV-2 infection on cells

7×10^4^ Vero E6 cells (ATCC #CRL-1586) were seeded into an Millicell EZ SLIDES chamber slides (Millipore) 24 hours before infection. On the day of infection, cells were pre-treated with 200 nM mitoxantrone or as a control with DMSO for 30 min. After 30 min treatment, Vero E6 cells were infected with live SARS-Cov-2 (USA-WA1/2020) at an MOI of 0.1 for 5 hours at 37°C, 5% CO2. Cells were then washed, fixed with 4% paraformaldehyde. The viral spike protein was stained with an in-house developed rabbit anti-spike antibody and goat anti-rabbit IgG (H+L) highly cross-adsorbed antibody with Alexa Fluor 488 (Invitrogen). Images were acquired using FluoView FV10i Confocal Laser Scanning Microscope (Olympus). All live virus infections were performed in a biosafety level 3 (BSL3) laboratory as approved by the Institutional Biosafety Committee (IBC).

### SARS-CoV-2 infection in a 3D EpiAirway model

Infection in the 3D EpiAirway model was carried out by University of Louisiana as a contracted service. In general, human tracheobronchial epithelial cells (EpiAirway™ from MatTek) were culture on inserts at ALI in 6-well plates. Prior to addition of drugs or virus, accumulated mucus from the tissue surface were removed by gently rinsing the apical surface twice with 400 μl TEER buffer. All fluids from the tissue surface were carefully removed to leave the apical surface exposed to the air. Mitoxantrone and control compounds were diluted into the assay medium (AIR-ASY-100). Tissues were pretreated with compounds for 1 hr on the apical and basal sides. After compound pretreatment, liquid was removed from the apical side, and virus (2019-nCoV/USA-WA1/2020 at MOI of 0.1) was inoculated in 0.15 mL assay medium onto the apical layer for 1 hour. Following 1 hour inoculation, virus-containing media was removed from apical layer and the apical side was washed with 400 μl TEER buffer. The basolateral medium was replaced with fresh maintenance medium. Media on the basolateral side was replaced with fresh compound containing maintenance medium at 24, 48, and 72hrs post-infection.

### TCID_50_ (50% Tissue Culture Infectious Dose) assay

Human bronchial epithelial cells (HBEC’s 3D-EpiAirway™) were seeded into culture inserts for 6-well plates one day before viral infection. Prior to addition of drugs or virus, accumulated mucus from the tissue surface were removed by gently rinsing the apical surface twice with 400 μl TEER buffer. All fluids from the tissue surface were carefully removed to leave the apical surface exposed to the air. Mitoxantrone was diluted into the assay medium and placed at room temperature before co-treatment with virus (MOI: 0.1) onto apical layer and basal layer for 1 hour. Following 1 hour treatment, virus was removed from apical layer and basolateral medium was replaced with fresh maintenance medium and compound at 24 hrs, 48 hrs and 72 hrs postinfection.

At 24 hours and 96 hours post-infection, the apical layer was washed with 0.4 mL of TEER buffer (PBS with Mg^2+^ and Ca^2+^) and aliquoted to separate microfuge tubes (1.5 mL). Eight-fold serial dilutions of apical layer supernatant sample concentrations were added to 96-well assay plates containing Vero E6 cells (20,000/well). The plates were incubated at 37 °C, 5% CO_2_ and 95% relative humidity. Following 3 days (72 ± 4 h) incubation, the plates were stained with crystal violet to measure cytopathic effect (CPE). Virus titers were calculated using the method of Reed and Muench (Reed et al., 1938). The TCID_50_ values were determined from duplicate samples.

### LDH Assay

Medium from the basolateral layer of the tissue culture inserts was removed 24- and 96-hours post-infection and diluted in LDH Storage Buffer as per the manufacturer’s instructions (LDH-Glo Cytotoxicity Assay, Promega). Samples (5 μl) were further diluted with LDH Buffer (95 μl) and incubated with an equal volume of LDH Detection Reagent. Luminescence was recorded after 60 minutes incubation at room temperature. A no cell control was included as a negative control to determine culture medium background and bleomycin included as a positive cytotoxic control.

### Gene expression and structural analyses

To study the Mitoxantrone’s effect on gene expression in HEK293T cells, we analyzed the mRNA sequencing data from our previous study, which is stored in the NCBI sequence read archive with accession ID: PRJNA645209. We used the web-based String protein interaction network program to analyze significantly up- or down-regulated genes. The identified networks were exported into Cytoscape for graph-making. The complex of Mitoxantrone with DNA-topoisomerase was analyzed by LigPlot ^34^ to reveal the interaction between Mitoxantrone and the enzyme.

To determine how Mitoxantrone interacts with DNA and topoisomerase, we used LigPlot to analyze the molecular interactions present in the published structure (PDB: 4g0v) ^27^.

### Image processing and statistical analyses

Confocal images were processed using the Zeiss Zen software. To measure fluorescence intensity, we used the open-source Fiji software. Images were converted to individual channels and regions of interest were drawn for measurement. Statistical analyses were performed using either Excel or GraphPad Prism 9. Data are presented as means□±□SEM, which was calculated by GraphPad Prism 9. *P* values were calculated by Student’s *t*-test using Excel. Nonlinear curve fitting and IC_50_ calculation was done with GraphPad Prism 9 using the inhibitor response three variable model or the exponential decay model. Images were prepared with Adobe Photoshop and assembled in Adobe Illustrator. Data processing and reporting are adherent to the community standards.

## Acknowledgments

The work was supported by the intramural research program of the National Institute of Diabetes, Digestive & Kidney Diseases (Y.Y.) and of the National Center for Advancing Translational Sciences (W.Z.) in the National Institutes of health, and by an FDA/CBER intramural SARS-CoV-2 fund (H.X.). The clinical isolate USA-WA1/2020 (ATCC# NR-5228) was deposited by the Centers for Disease Control and Prevention and obtained through BEI Resources, NIAID, NIH: SARS-Related Coronavirus 2.

## Author contributions

Ye, Y., Zhang, Q., Xie, H., Zheng, W., Chen. C. planned the study. Zhang, Q., Radvak, P., Lee, J., Xu, M. conducted the experiments and analyzed the data, Xu, Y. analyzed the Mitoxantrone-DNA topoisomerase structure, Cao-dao, V. analyzed the gene expression data, Zhang, Q. and Ye Y. wrote the paper.

